# Sequence and epigenetic landscapes of active and silenced nucleolus organizers in Arabidopsis

**DOI:** 10.1101/2023.06.07.544131

**Authors:** Dalen Fultz, Anastasia McKinlay, Ramya Enganti, Craig S. Pikaard

**Affiliations:** Howard Hughes Medical Institute, Indiana University; Bloomington, IN, USA; Department of Biology, Indiana University; Bloomington, IN, USA; Department of Molecular and Cellular Biochemistry, Indiana University; Bloomington, IN, USA

## Abstract

*Arabidopsis thaliana* has two ribosomal RNA gene loci, nucleolus organizer regions *NOR2* and *NOR4*, whose complete sequences remain undefined. Ultra-long DNA sequences assembled using an unconventional approach yielded 5.5 and 3.9 Mbp sequences for *NOR2* and *NOR4* (in the reference strain, Col-0), revealing their distinct gene subtype compositions. RNA sequencing and identification of genes associated with flow-sorted nucleoli of wild-type or silencing-defective mutant plants shows that most of *NOR4* is comprised of active genes whereas most, but not all, *NOR2* genes are epigenetically silenced. Long intervals of low CG and CHG methylation overlap regions of gene activity and gene subtype homogenization. Collectively, the data reveal the genetic and epigenetic landscapes of the NORs and implicate transcription in rRNA gene concerted evolution.

**One Sentence Summary:** *NOR2* and *NOR4* sequences fill genome gaps and enable megabase-scale analyses of rRNA gene regulation and concerted evolution.

## Main Text

In eukaryotes, ribosomal RNA (rRNA) genes transcribed by RNA Polymerase I (Pol I) are repeated in long tandem arrays, typically spanning millions of basepairs (*1, 2*). These chromosomal loci are known as nucleolus organizer regions (NORs) because the nucleolus, the most prominent substructure of the nucleus, forms around rRNA genes that are transcribed and processed into 18S, 25-28S and 5.8S catalytic rRNAs (*3, 4*). The rRNAs are then assembled with ∼80 ribosomal proteins and 5S rRNA to form ribosomes, the protein-synthesizing machines of cells (*5-7*).

rRNA gene repeats within a species are extremely similar in sequence complexity yet can be differentially expressed, accounting for the phenomenon known as nucleolar dominance (*8-10*). This epigenetic phenomenon was initially discovered in genetic hybrids and describes the preferential activity of NORs inherited from one progenitor. However, nucleolar dominance is not unique to hybrids, also occurring in pure species (non-hybrids) to regulate the number of active rRNA gene repeats (*9, 11, 12*). In *Arabidopsis thaliana* Col-0, ∼50% of the rRNA genes are silenced during early post-embryonic development (*13*) in a process involving DNA cytosine methylation and histone deacetylation (*14, 15*). Genetic segregation analyses of a small set of differentially expressed rRNA gene subtypes found that silenced subtypes mapped to the NOR on chromosome 2, *NOR2* whereas active subtypes mapped to *NOR4* (*16, 17*). However, in a Col-0 line in which a large portion of *NOR4* was replaced by sequences of *NOR2*, the transposed *NOR2* genes escaped silencing (*18*). Collectively, these results suggest that rRNA gene activity is not dictated by individual gene sequences but, instead, somehow depends on NOR affiliation, highlighting a need to determine the sequences and functional organizations of the NORs (*19, 20*).

### Variable-length elements define at least 74 rRNA gene subtypes

At *NOR2* and *NOR4*, rRNA genes are repeated head-to-tail, oriented such that transcription occurs towards the centromere (Fig. 1A) (*21, 22*). Each transcription unit is separated by an intergenic spacer (IGS) that includes the gene promoter, 0-4 spacer promoters (SPs)(*23*), and variable numbers of short repeats bearing *Sal* I restriction endonuclease sites (*24*). Sequence elements within the transcribed region include a 5’ External Transcribed Spacer (5’ ETS), 18S, 5.8S and 25S catalytic rRNAs, two Internal Transcribed Sequences, ITS1 and ITS2, and a 3’ ETS. Gene promoter, rRNA and ITS sequences are nearly identical among rRNA gene repeats, except for rare deletions (Fig. 1B) (*22*). However, substantial variation occurs within IGS and ETS regions due to variable numbers of Sal repeats, spacer promoters, 5’ ETS repeats, or 3’ ETS repeats, making each of these elements variable in length (Fig. 1B). Using Pacific Bioscience SMRT sequencing of ∼10 kbp rRNA gene units cut from genomic DNA by I-*Ppo*I (fig. S1), we expanded upon the known variation (*22, 24, 25*), doubling the number of variable-length elements (VLEs). The 59 VLEs summarized in Fig. 1B occur in 74 permutations, thus defining a minimum of 74 rRNA gene subtypes (fig. S2A).

**Figure 1.**
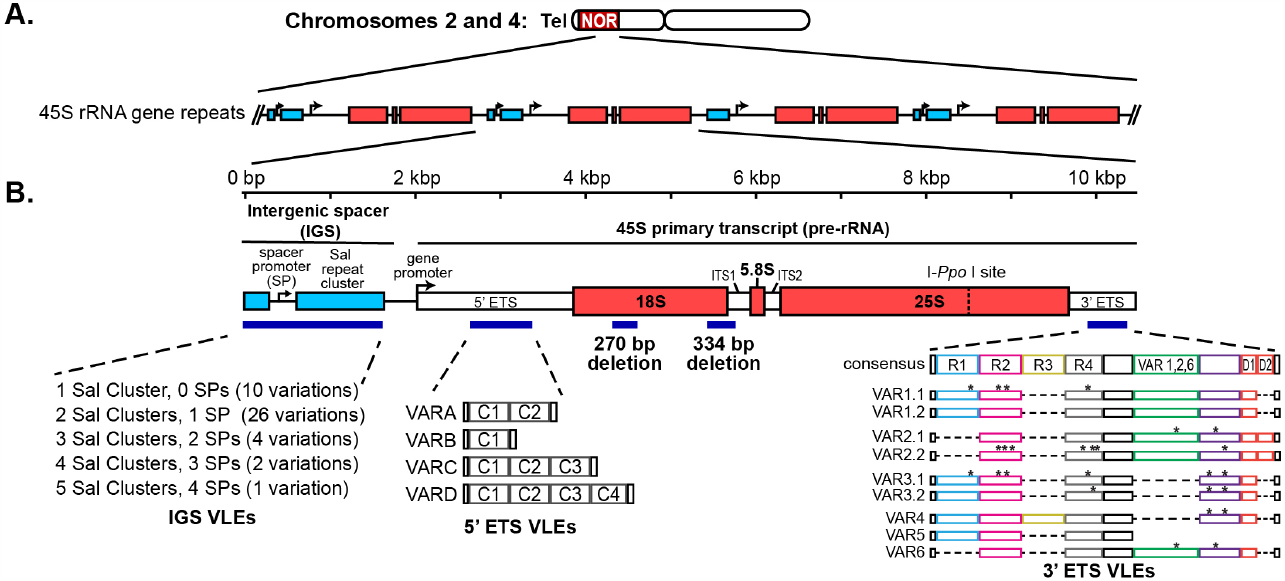
Organization of the *Arabidopsis thaliana* NORs. (**A**) The NORs abut the telomeres on the short arms of chromosomes 2 and 4 and consist of tandemly repeated copies of 45S rRNA genes. (**B**) Diagram of a single rRNA gene repeat, showing the locations of Variable Length Elements (VLEs). “C” repeats in the 5’ ETS are 310 bp in length. Asterisks in the 3’ETS “R” repeats and elsewhere denote single nucleotide polymorphisms relative to a consensus reference sequence.

### Sequencing, assembly, and compositional analyses of NOR2 and NOR4

Sequence conservation among rRNA genes precludes their discrimination using short-read DNA sequencing, making NORs notoriously difficult loci to assemble. However, we hypothesized that rare VLE patterns spanning multiple genes might provide landmarks for NOR assembly. To this end, we performed Oxford Nanopore Technology (ONT) sequencing of *A. thaliana* genomic DNA, obtaining reads averaging ∼40-50 kbp but including numerous reads longer than 200 kbp (table S1). Whole genome assembly using the long ONT reads recapitulated the TAIR10 genome assembly (fig. S3), confirming comprehensive genome coverage within the read set. However, neither NOR was substantially assembled, as is true for the TAIR10 reference genome assembly and several recent long-read assemblies (*26-28*).

To fully assemble the NORs, we used two complementary approaches. In one approach, ONT reads containing rRNA gene sequences were compared to a single rRNA gene reference sequence, namely the sequence of gene subtype #10 (see file S1 and fig. S2) to generate two-dimensional similarity matrices (dot-plots). Individual dot-plots displaying distinct and rare patterns were then manually assembled, like puzzle pieces, yielding longer contigs, as illustrated in Fig. 2A. The second approach involved a computational NOR assembly pipeline (Fig. 2B).

**Figure 2.**
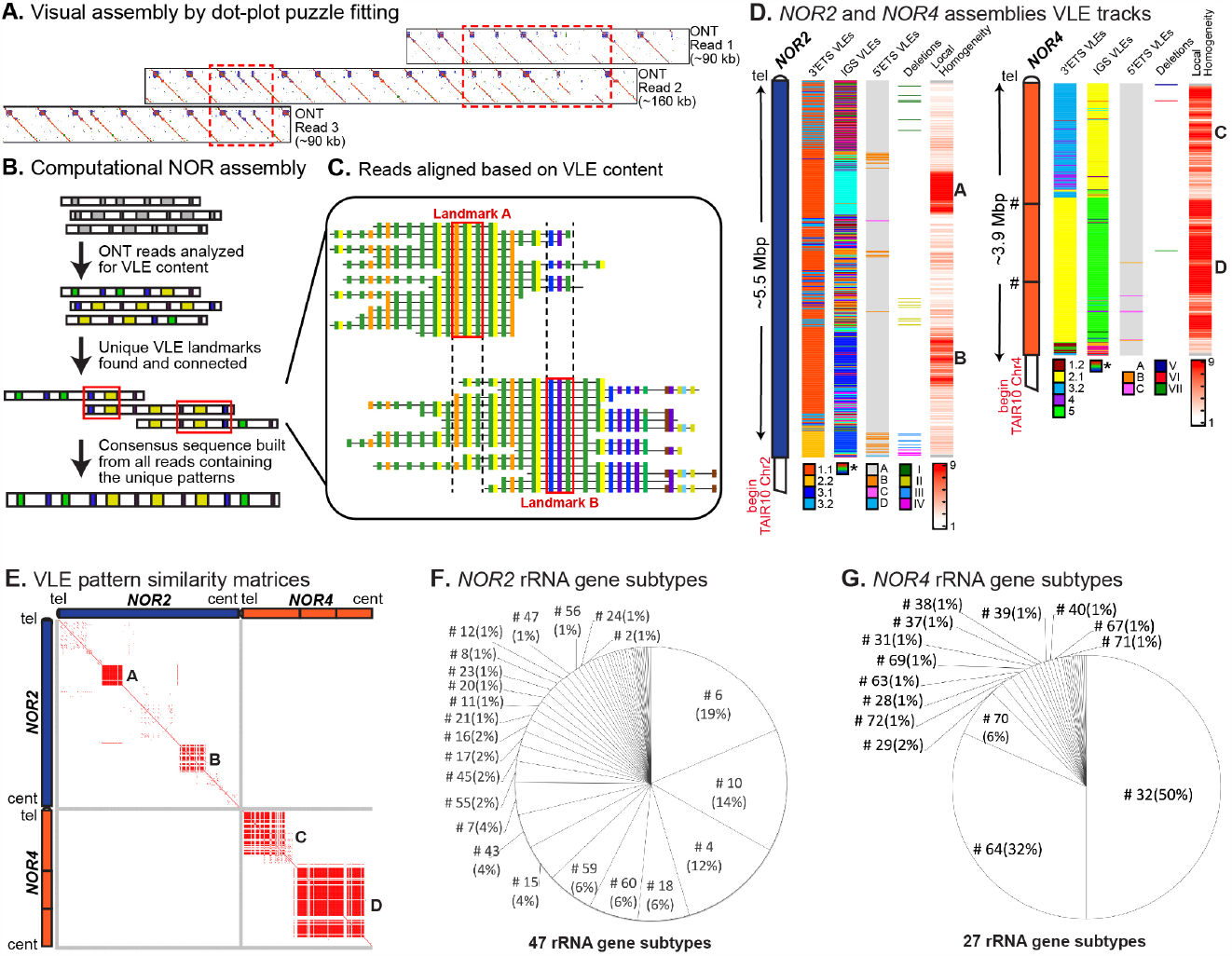
Assembly of *NOR2* and *NOR4*. (**A**) Example of an assembly of overlapping dot-plots. (**B**) Schematic of the computational assembly pipeline. (**C**) Diagram depicting aligned reads at two adjacent VLE landmarks. (**D**). Graphical summary of the NOR assemblies, with tracks for distinct VLEs, partial gene deletions, and local homogeneity (number of identical gene subtypes within a nine gene sliding window). Color codes are provided below each track, except for IGS VLEs which have too many types and associated colors (forty-three) to provide a meaningful color code. (**E**) Similarity matrices comparing VLE content within and between NORs. Each red point represents a perfect match of VLEs, within a three gene window, in the sequences being compared. (**F-G**). Pie charts representing the relative abundance of gene subtypes within *NOR2* or *NOR4*. See fig. S2 for subtype details.

The pipeline first determined the order of VLEs within long (>85 kbp) rRNA gene-containing ONT reads that collectively provided ∼100X coverage for an estimated ∼750-800 rRNA genes per haploid genome (*29*) (see methods). We then identified sets of VLEs predicted to be unique based on their frequency within the read set (file S2) and used them as landmarks for aligning and grouping overlapping ONT reads, ultimately connecting the landmarks (see Fig. 2C and files S3-S7) to yield full-length sequence assemblies of 5.5 Mbp and 3.9 Mbp for *NOR2* and *NOR4*, respectively (Fig. 2D). Multiple sequence alignment of overlapping reads enabled consensus sequence polishing for the two NORs (see Methods), with comparisons to 18S and 25S rRNA sequences, which are essentially invariant (*17*), indicating that the final *NOR2* and *NOR4* sequences have a basepair accuracy of ∼99.85% (file S8). The polished NOR sequences were deposited at Genbank (BankIt2506918 and BankIt2708174).

We quality checked the *NOR2* and *NOR4* assemblies in three ways. First, we compared the frequency of VLE patterns in ONT reads versus the NOR assemblies, obtaining a correlation value of R^2^ = 0.985 (file S9.) This indicates that the assemblies reflect the VLE patterns in the raw data. Second, ONT reads of computationally assembled contigs were visually examined using the dot-plot puzzle-fitting approach. Third, we used custom-designed CRISPR guide RNAs complementary to alternative 3’ ETS VLEs to catalyze Cas9 cutting of genomic DNA. We then separated the DNA fragments by contour-clamped homogeneous electric field (CHEF) electrophoresis and performed Southern blotting with a rDNA probe (fig S4). The Cas9 digestion products were consistent with those predicted from the NOR assemblies.

*NOR2* has 518 rRNA gene repeats, based on the count of 25S rRNA sequences, whereas *NOR4* has 376 genes. Fig. 2D shows their organizations with respect to 3’ETS, IGS and 5’ETS VLEs, or internal deletions, each displayed as a vertical track. Another track depicts the number of times any single rRNA gene subtype is repeated within a sliding window spanning 9 genes (∼90 kbp), providing a measure of local subtype homogeneity. Subtype content and distribution patterns are unique to each NOR, as illustrated in similarity matrices comparing VLE features within and between the NORs (Fig. 2E).

Some VLEs are shared by genes at both NORs but none of the 74 rRNA gene subtypes are common to both NORs (fig. S2B, fig. S5). *NOR2* is comprised of 47 rRNA gene subtypes (Figs. 2F, 2G, 3A, and 3B), mostly VAR1.1-class genes (subtypes 1-26 in Fig. 3A and fig. S2A) interspersed with VAR3 genes of both the 3.1 (subtypes 47-53) and 3.2 subclasses (subtypes 54-70). Two regions of high local subtype homogeneity account for ∼25% of *NOR2* (Figs 2D and 2E). Homogenous region A is enriched for gene subtype 6 (Fig. 3A), the most abundant *NOR2* subtype, comprising 19% of the NOR (Fig. 2F). Region B is enriched for subtype 10, the second most abundant *NOR2* subtype (14%; Figs. 2F and 3A).

**Figure 3.**
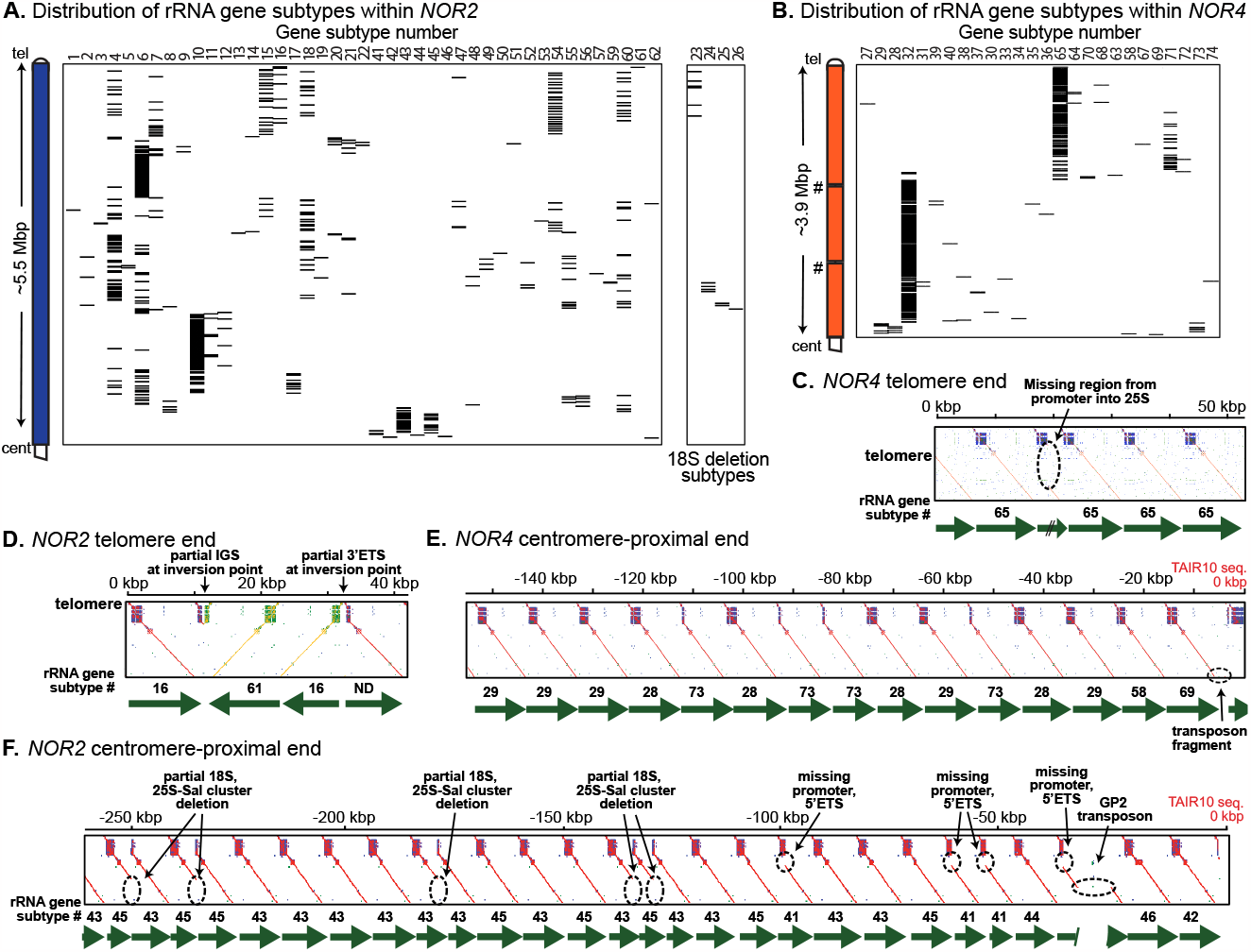
Structural features of *NOR2* and *NOR4*. (**A-B**) Positions of individual gene subtypes throughout *NOR2* and *NOR4*. The group separated at the right of panel A are subtypes with deletions in 18S rRNA sequences (**C-F**) Similarity matrices for the telomere and centromere-proximal ends of *NOR2* and *NOR4*. Distinctive features are highlighted with dashed circles. Numbering is relative to the telomere-rRNA gene junction in panels (C) and (D) and relative to the start of the TAIR10 chromosome 2 and 4 assemblies for (E) and (F).

*NOR4* is comprised of 27 rRNA gene subtypes (Fig. 2G, figS2B), with three accounting for 88% of the NOR (Fig. 2G) and clustering within homogenous regions C and D (Figs. 2D-E). Region C is composed of VAR3.2-class genes, primarily subtype 65 genes that account for 32% of *NOR4* (Figs 2G and 3B), interspersed with VAR4-class genes, primarily subtype 71 genes comprising 6% of the NOR (Figs 2G and 3B). Region D consists primarily of VAR2.1 class genes, specifically subtype 32, which comprises 50% of *NOR4* (Figs 2G and 3B).

At two positions within *NOR4*, marked by ‘#’ symbols in Fig. 2D (and subsequent figures), the contigs on either side end in long stretches of subtype 32 repeats and cannot be joined unambiguously at specific positions. Comparison of subtype 32 abundance in raw sequencing reads versus the *NOR4* assembly suggests that ∼31 copies of subtype 32 may be unaccounted for in the assembly (file S9). If so, the additional genes would be present at one or both locations indicated by the # symbols, potentially making *NOR4* somewhat longer than indicated.

The telomere and centromere-proximal ends of *NOR2* and *NOR4* have distinct structural features (Figs 3C-F). At the telomeric end of *NOR4*, the third gene has an internal deletion of several kbp (Fig. 3C), one of only four structural anomalies within the entirety of *NOR4*. The other anomalies are two genes with 25S deletions (Fig. 2D, ‘Deletions’ track) and one gene, near the centromere-proximal end, disrupted by a GP2 (Gypsy family) transposon (Fig. 3E). By contrast, *NOR2* has numerous genes with structural anomalies. At the telomere-proximal end, an inversion of ∼20 kbp occurred (Fig. 3D). At the centromere-proximal end, more than one-third of the genes in the final 250 kbp have internal deletions, and a GP2 transposon interrupts the third gene from the end (Fig. 3F). In central regions of *NOR2*, multiple genes have internal deletions. These deletions were previously noted among rRNA genes cloned within bacterial artificial chromosomes (*22*). In total, *NOR2* has ∼20 genes whose internal deletions would likely render them non-functional (see Fig. 2D ‘Deletions’ track).

### Locations of active and silenced rRNA gene subtypes within NOR2 and NOR4

To determine where active rRNA genes are located within the NORs, we first identified informative nucleotide polymorphisms using Illumina whole genome sequencing, conducted at a depth of coverage of ∼163X (fig. S6A and file S10), and determined the positions of these polymorphisms in our NOR assemblies (fig. S6B). In parallel, we performed Illumina sequencing of reverse-transcribed leaf RNA and identified polymorphic nucleotides within this read set. Thirteen of forty-two nucleotide polymorphisms specific to *NOR4* genes were detected by RNA-seq. Some are common to genes broadly distributed throughout the NOR, any of which could be responsible for the transcripts detected. Others localize to specific regions. Collectively, the broad and region-specific markers indicate that genes in the central region of *NOR4* are highly expressed. At the telomeric and centromere-proximal ends of *NOR4*, the polymorphic markers specific to these regions are not detected in the RNA, suggesting that transcriptional activity wanes near the NOR ends (Figure 4A).

**Figure 4.**
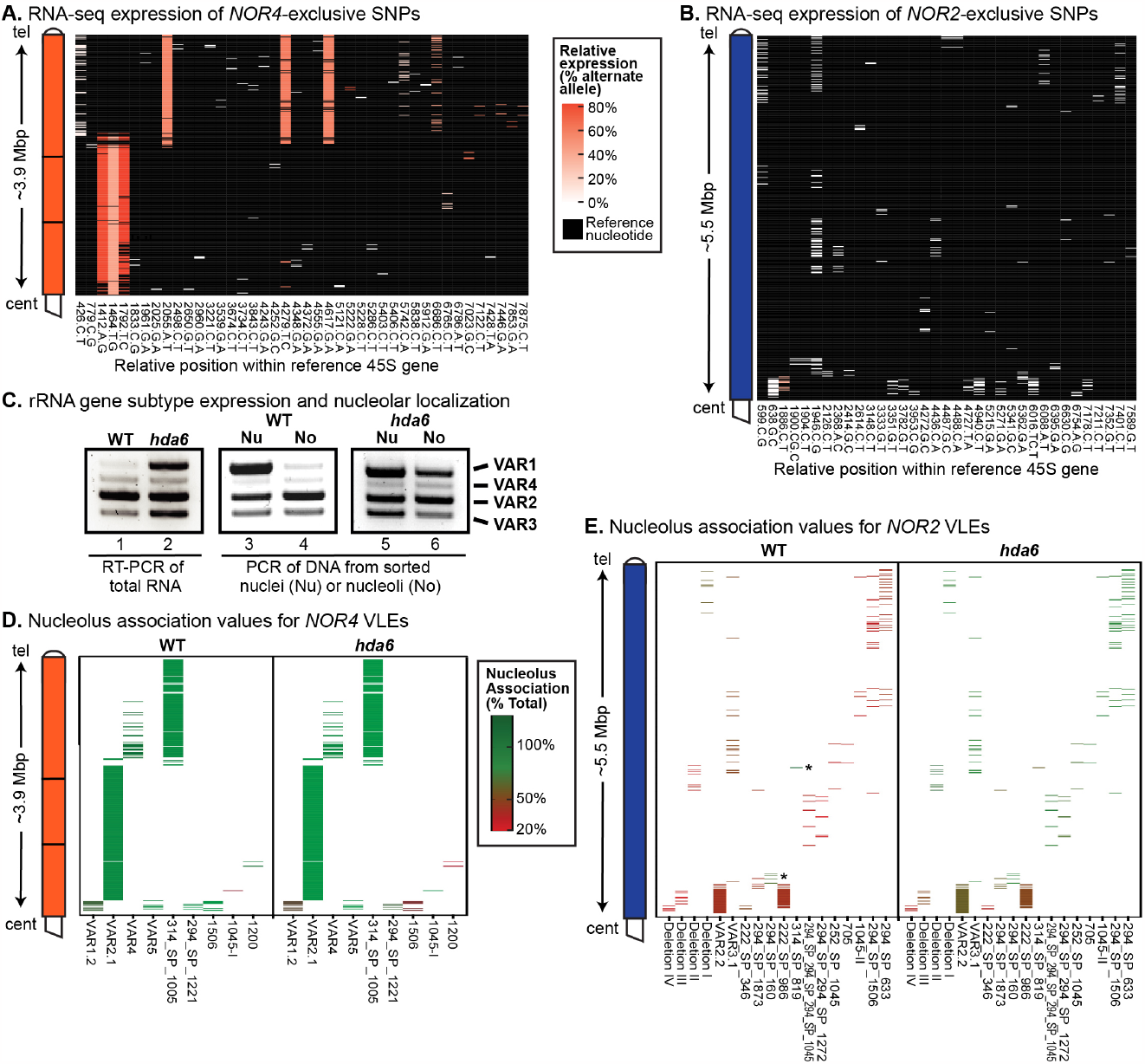
rRNA gene activity and nucleolar association maps for *NOR2* and *NOR4*. (**A-B**) Heatmaps of expression levels for *NOR4* or *NOR2*-specific SNPs, each represented by a horizontal line on the y-axis. SNP names on the X axis include the position (number) in the ribosomal gene reference sequence followed by the reference and alternate nucleotide. White to red colors indicate level of expression based on normalized RNA-seq read counts. Black indicates reference nucleotide positions with no alternative SNPs. (**C**) RT-PCR of the 3’ETS VAR region, comparing WT and *hda6* mutants (left panel) and comparing nuclear versus nucleolus-associated genes (middle and right panel). (**D-E**) Heatmaps depicting degree of nucleolus association for *NOR4* or *NOR2*-specific VLEs. Asterisks denote rare *NOR2* VLEs that are nucleolus associated. Experiments were performed using leaf tissue of mature plants.

Thirty-five polymorphisms were *NOR2*-specific, representing genes distributed throughout the NOR (Fig. 4B). Only one was detected among expressed RNAs, C1886T, present within a cluster of genes near the centromere-proximal end, indicating that *NOR2* is mostly, but not entirely, inactive.

### rRNA genes distributed throughout NOR4 co-purify with nucleoli

The abundant nucleolar protein, Fibrillarin, when expressed as a fusion protein with YFP, enables the fluorescence-activated flow-sorting of nuclei liberated by cell disruption or of nucleoli that persist upon disruption of nuclei by sonication (*15*). Using a PCR primer pair that flanks the 3’ ETS variable region, amplification products of genes bearing the VAR1, 2, 3 or 4 VLEs (*16*) are all detected in the DNA of nuclei (Fig. 4C lanes 3 and 5). However, VAR1-class genes are depleted in isolated nucleoli (Fig. 4C, lane 4), reflecting their selective silencing (Fig 4C, lane 1). However, if silencing is prevented in an *hda6* mutant, VAR1 genes are expressed at high levels (Fig. 4C, lane 2) (*14, 15*) and co-purify with nucleoli (Fig. 4C, lane 6). These experiments show that nucleolar localization can be used as a proxy for rRNA gene transcriptional activity. Importantly, this allows active genes to be identified based on VLEs in the DNA, thereby allowing discrimination among rRNA gene subtypes whose transcripts are indistinguishable by RNA-seq.

Using ONT sequencing of DNA, we identified the rRNA gene subtypes that co-purify with flow-sorted nucleoli or whole nuclei (as controls). The ONT reads averaged ∼3 kbp in length (table S2), sufficient for VLE identification. The abundance of 3’ETS VLE reads from nuclear DNA matched their abundance in the NOR assemblies (fig S7), indicating that all subtypes are detected without apparent bias. Next, we calculated the level of nucleolar enrichment for NOR-specific VLEs. *NOR4* rRNA genes are enriched in nucleoli, as demonstrated for nine *NOR4*-specific VLEs (Fig. 4D), two of which (VAR2.1 and 314_SP_1005) represent homogenous regions C and D (see Figs. 2D-E). Nucleolar association of the *NOR4* VLEs was mostly unchanged in an *hda6* mutant versus wild-type except for decreased expression of genes near the centromere-proximal end of the NOR. By contrast, dramatic differences were apparent for *NOR2 –*specific VLEs in wild-type versus *hda6* (Fig. 4E). In wild-type plants, 16 of the 18 VLEs examined, distributed throughout *NOR2*, are depleted in nucleoli relative to nuclei but become nucleolus-enriched in *hda6* plants, indicative of HDA6-dependent silencing. Notably, genes near the centromere-proximal end of *NOR2* failed to become nucleolus-enriched in *hda6* plants. Many genes in this region are missing promoters (see Fig. 3F), likely precluding their participation in nucleolus formation. Two of the eighteen *NOR2*-specific VLEs were nucleolus-associated (see asterisks in Fig. 4E), even in wild-type, with one present in a small gene cluster near a cluster also identified by RNA-seq (Fig. 4B), consistent with pockets of gene expression within *NOR2*.

### Cytosine methylation landscapes of NOR2 and NOR4

We examined CG, CHG and CHH methylation throughout *NOR2* and *NOR4* using ONT methylation calling tools (*30*), calculating the percent methylation within 500 bp sliding windows (Figs 5A,B). Most of *NOR2* has extremely high CG methylation, moderate CHG methylation and relatively low CHH methylation (Figs 5A). An exception is *NOR2* homogenous region B, which is hypomethylated in all three contexts (Fig. 5A and fig. S8). At *NOR4*, a central region of ∼ 2 Mbp has low methylation levels, in all contexts, but is flanked, on both sides, by ∼1 Mbp intervals of high methylation (Fig. 5D). Zooming in shows that within regions of high CG methylation, methylation is high across entire gene units except near the gene promoter, which is hypomethylated in most, but not all, genes (fig. S8). Conversely, in low methylation regions, the hypomethylation is uniform except in the 3’ETS region, where a peak of methylation occurs in most gene repeats (fig. S8). The most highly methylated regions of *NOR2* and *NOR4* correlate with regions of low or undetectable gene expression (Figs. 5A-B, RNA track). Conversely, the hypomethylated central region of *NOR4* overlaps the positions of actively transcribed gene subtypes (Fig. 5B, RNA track).

**Figure 5.**
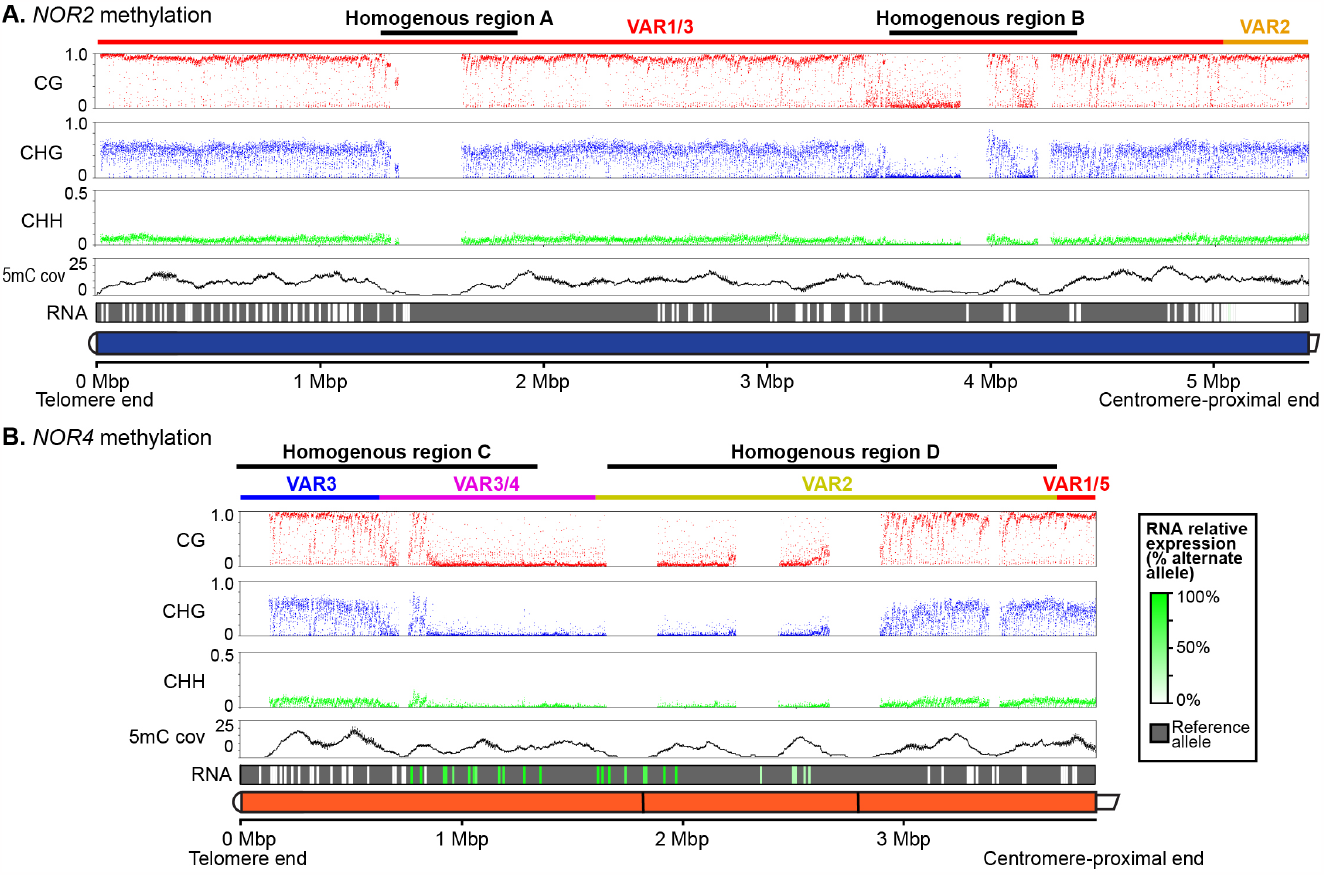
Methylation landscapes of *NOR2* and *NOR4*. (**A-B**) 5mC frequencies in the CG, CHG, and CHH contexts at *NOR2* and *NOR4*. Each colored point represents the frequency calculated within a 500 bp sliding window, with 250 bp steps. Homogenous regions where there are no data points reflect a paucity of uniquely mapped reads needed to surpass the threshold for 5mC calling. The 5mC cov (coverage) track shows the number of reads that were retained by the methylation calling pipeline. The RNA track shows the relative expression levels of SNPs, colored from white (not expressed) to shades of green (expressed). The gray background corresponds to nucleotides of the reference sequence. The DNA sequenced was from inflorescence apex tissue.

## Discussion

Using an unconventional sequence assembly strategy, our determination of the *NOR2* and *NOR4* sequences, from their telomere-capped ends to their centromere-proximal ends, fills the last major gaps that remain in the Arabidopsis genome following the determination of the centromeric regions (*26*). The two NOR sequences add 9.4 Mbp of new genome sequence information, equivalent to ∼6.5 % of the genome.

A surprising finding is that none of the 74 gene subtypes defined by VLE content is common to both NORs. Instead, the NOR have unique sets of rRNA gene subtypes, allowing subtype-specific expression to be mapped to specific positions within *NOR2* or *NOR4*.

Collectively, the data show that genes distributed throughout the 5.5 Mbp of *NOR2* are inactive, with only rare exceptions, whereas genes throughout a large central region of *NOR4* are active. Inactive *NOR2*-specific gene subtypes are epigenetically silenced, in an *HDA6*-dependent manner (Fig. 4).

Prior studies showed that silencing of rRNA genes with VAR1-class 3’ ETS sequences, which primarily map to NOR2 (*16*) requires the DNA methyltransferases MET1 and CMT2 in addition to HDA6 and Histone H3 lysine 9 methyltransferases (*13, 15, 31, 32*). Another study showed that histone modification and DNA methylation occur in a concerted manner at both silenced and active rRNA genes (*33*). These findings provide context for the ∼2 Mbp hypomethylated region at the center of *NOR4*, which overlaps the positions of rRNA gene subtypes that are transcribed and associated with the nucleolus. Our interpretation is that this hypomethylated region is the likely source of most rRNAs synthesized in the leaf and inflorescence tissues used in our study. Our observation that *NOR4* subtype expression is mostly unchanged in *hda6*, relative to wild-type plants, suggests that lower gene expression at the telomere and centromere-proximal ends of *NOR4* is distinct from *NOR2* gene silencing.

Regions of high gene activity and/or low cytosine methylation within *NOR2* and *NOR4* correlate with regions of high gene subtype homogeneity. Concerted evolution of rRNA genes (*34, 35*) is thought to occur via gene conversion, or recombination following slipped mispairing of NOR sequences (*36*). Our results suggest that transcription may facilitate local rRNA gene homogenization via one or both mechanisms. Consistent with this hypothesis, studies in *Saccharomyces cerevisiae* showed that mitotic recombination and gene conversion events are stimulated by rRNA gene promoter sequences (*37-39*).

The lack of gene subtypes common to both *NOR2* and *NOR4*, but the occurrence of shared VLEs (see fig. S5), suggests that co-evolution of the genes within the two NORs occurs through events involving sequence intervals shorter than complete gene units. The correlation between hotspots of transcription and hotspots of gene subtype homogenization is consistent with transcription acting as a driver of homogenization, perhaps explaining how two gene subtypes of *NOR4* have come to occupy 82% of the NOR. Conversely, the substantial heterogeneity *NOR2* (47 subtypes versus 27 at *NOR4*), including the persistence of genes with debilitating structural anomalies, may reflect its reduced transcriptional activity.

## References and Notes

## Acknowledgments

We thank the Indiana University Bloomington (IUB) Center for Genomics and Bioinformatics and the IUB Flow Cytometry Core Facility for expert assistance, the IUB Pervasive Technology Institute for computing resources and Dr. Feng Wang for the Cas9 guide RNA design script. This article is subject to HHMI’s Open Access to Publications policy. HHMI lab heads have previously granted a nonexclusive CC BY 4.0 license to the public and a sublicensable license to HHMI in their research articles. Pursuant to those licenses, the author-accepted manuscript of this article can be made freely available under a CC BY 4.0 license immediately upon publication.

## Author contributions

Conceptualization: DF, AM, CSP

Data curation: DF, AM

Methodology: DF, AM, RE, CSP

Investigation: DF, AM, RE

Visualization: DF, AM, RE

Funding acquisition: CSP

Project administration: DF, AM, RE, CSP

Software: DF

Supervision: CSP

Writing – original draft: DF, AM, CSP

Writing – review & editing: DF, AM, RE, CSP

## Data and materials availability

NOR sequences have been deposited at GenBank (BankIt2506918 and BankIt2708174). PacBio, ONT, and Illumina sequencing datasets have been deposited at NCBI (BioProject PRJNA982841). Custom scripts for NOR assembly steps have been submitted to GitHub. All sequences, datasets and computational tools will become publicly available upon publication.

## Supplementary Materials

Materials and Methods

Figs. S1 to S8

Tables S1 to S2

Data files S1 to S10

## References

1. E. O. Long, I. B. Dawid, Repeated genes in eukaryotes. Annu Rev Biochem 49, 727–764 (1980).

2. S. A. Gerbi, in Molecular Evolutionary Genetics, R. J. McIntyre, Ed. (Plenum Press, New York, 1985), pp. 419–517.

3. F. M. Boisvert, S. van Koningsbruggen, J. Navascues, A. I. Lamond, The multifunctional nucleolus. Nat Rev Mol Cell Biol 8, 574–585 (2007).

4. T. Pederson, The nucleolus. Cold Spring Harb Perspect Biol 3, DOI 10.1101/cshperspect.a000638 (2011).

5. T. W. Turowski, D. Tollervey, Cotranscriptional events in eukaryotic ribosome synthesis. Wiley interdisciplinary reviews. RNA 6, 129–139 (2015).

6. J. Bassler, E. Hurt, Eukaryotic Ribosome Assembly. Annu Rev Biochem 88, 281–306 (2019).

7. J. Saez-Vasquez, M. Delseny, Ribosome Biogenesis in Plants: From Functional 45S Ribosomal DNA Organization to Ribosome Assembly Factors. Plant Cell 31, 1945–1967 (2019).

8. R. H. Reeder, Mechanisms of nucleolar dominance in animals and plants. J. Cell Biol. 101, 2013–2016 (1985).

9. B. McStay, Nucleolar dominance: a model for rRNA gene silencing. Genes Dev 20, 1207–1214 (2006).

10. C. S. Pikaard, Nucleolar dominance and silencing of transcription. Trends Plant Sci. 4, 478–483 (1999).

11. N. Borowska-Zuchowska et al., Switch them off or not: selective rRNA gene repression in grasses. Trends Plant Sci DOI 10.1016/j.tplants.2023.01.002, (2023).

12. S. Tucker, A. Vitins, C. S. Pikaard, Nucleolar dominance and ribosomal RNA gene silencing. Curr Opin Cell Biol 22, 351–356 (2010).

13. K. W. Earley et al., Mechanisms of HDA6-mediated rRNA gene silencing: suppression of intergenic Pol II transcription and differential effects on maintenance versus siRNA-directed cytosine methylation. Genes Dev 24, 1119–1132 (2010).

14. K. Earley et al., Erasure of histone acetylation by Arabidopsis HDA6 mediates large-scale gene silencing in nucleolar dominance. Genes Dev 20, 1283–1293 (2006).

15. F. Pontvianne et al., Subnuclear partitioning of rRNA genes between the nucleolus and nucleoplasm reflects alternative epiallelic states. Genes and Development 27, 1545–1550 (2013).

16. C. Chandrasekhara, G. Mohannath, T. Blevins, F. Pontvianne, C. S. Pikaard, Chromosome-specific NOR inactivation explains selective rRNA gene silencing and dosage control in Arabidopsis. Genes Dev 30, 177–190 (2016).

17. F. A. Rabanal et al., Epistatic and allelic interactions control expression of ribosomal RNA gene clusters in Arabidopsis thaliana. Genome Biol 18, DOI 10.1186/s13059-13017-11209-z (2017).

18. G. Mohannath, F. Pontvianne, C. S. Pikaard, Selective nucleolus organizer inactivation in Arabidopsis is a chromosome position-effect phenomenon. Proc Natl Acad Sci U S A 113, 13426–13431 (2016).

19. C. S. Pikaard, C. Chandrasekhara, A. McKinlay, R. Enganti, D. Fultz, Reaching for the off switch in nucleolar dominance. Plant J, (2023).

20. B. McStay, Nucleolar organizer regions: genomic ‘dark matter’ requiring illumination. Genes Dev 30, 1598–1610 (2016).

21. G. P. Copenhaver, C. S. Pikaard, RFLP and physical mapping with an rDNA-specific endonuclease reveals that nucleolus organizer regions of Arabidopsis thaliana adjoin the telomeres on chromosomes 2 and 4. Plant J 9, 259–272 (1996).

22. J. Sims, G. Sestini, C. Elgert, A. von Haeseler, P. Schlogelhofer, Sequencing of the Arabidopsis NOR2 reveals its distinct organization and tissue-specific rRNA ribosomal variants. Nat Commun 12, DOI 10.1038/s41467-41020-20728-41466 (2021).

23. J. H. Doelling, R. J. Gaudino, C. S. Pikaard, Functional analysis of Arabidopsis thaliana rRNA gene and spacer promoters in vivo and by transient expression. Proc Natl Acad Sci U S A 90, 7528–7532 (1993).

24. P. Gruendler, I. Unfried, K. Pascher, D. Schweizer, rDNA intergenic region from Arabidopsis thaliana. Structural analysis, intraspecific variation and functional implications. J Mol Biol 221, 1209–1222 (1991).

25. K. Havlova et al., Variation of 45S rDNA intergenic spacers in Arabidopsis thaliana. Plant Mol Biol, (2016).

26. M. Naish et al., The genetic and epigenetic landscape of the Arabidopsis centromeres. Science 374, eabi7489 (2021).

27. X. Hou, D. Wang, Z. Cheng, Y. Wang, Y. Jiao, A near-complete assembly of an Arabidopsis thaliana genome. Mol Plant 15, 1247–1250 (2022).

28. B. Wang et al., High-quality Arabidopsis thaliana Genome Assembly with Nanopore and HiFi Long Reads. Genomics, proteomics & bioinformatics / Beijing Genomics Institute 20, 4–13 (2022).

29. G. P. Copenhaver, C. S. Pikaard, Two-dimensional RFLP analyses reveal megabase-sized clusters of rRNA gene variants in Arabidopsis thaliana, suggesting local spreading of variants as the mode for gene homogenization during concerted evolution. Plant J 9, 273–282 (1996).

30. P. Ni et al., Genome-wide detection of cytosine methylations in plant from Nanopore data using deep learning. Nat Commun 12, 5976 (2021).

31. F. Pontvianne et al., Histone methyltransferases regulating rRNA gene dose and dosage control in Arabidopsis. Genes Dev 26, 945–957 (2012).

32. G. Mohannath et al., DNA hypermethylation and condensed chromatin correlate with chromosome-specific rRNA gene silencing in Arabidopsis. BioRxiv, (2023).

33. R. J. Lawrence et al., A concerted DNA methylation/histone methylation switch regulates rRNA gene dosage control and nucleolar dominance. Mol Cell 13, 599–609 (2004).

34. G. Dover et al., in Genome evolution, G. A. Dover, R. B. Flavell, Eds. (Academic Press, London, 1982), pp. 343–372.

35. E. A. Zimmer, S. L. Martin, S. M. Beverley, Y. W. Kan, A. C. Wilson, Rapid duplication and loss of genes coding for the alpha chains of hemoglobin. Proc Natl Acad Sci U S A 77, 2158–2162 (1980).

36. D. Liao, Concerted evolution: molecular mechanism and biological implications. Am J Hum Genet 64, 24–30 (1999).

37. S. E. Stewart, G. S. Roeder, Transcription by RNA polymerase I stimulates mitotic recombination in Saccharomyces cerevisiae. Mol Cell Biol 9, 3464–3472 (1989).

38. K. Voelkel-Meiman, R. L. Keil, G. S. Roeder, Recombination-stimulating sequences in yeast ribosomal DNA correspond to sequences regulating transcription by RNA polymerase I. Cell 48, 1071–1079 (1987).

39. K. Voelkel-Meiman, G. S. Roeder, Gene conversion tracts stimulated by HOT1-promoted transcription are long and continuous. Genetics 126, 851–867 (1990).

40. F. Barneche, F. Steinmetz, M. Echeverria, Fibrillarin genes encode both a conserved nucleolar protein and a novel small nucleolar RNA involved in ribosomal RNA methylation in Arabidopsis thaliana. J Biol Chem 275, 27212–27220 (2000).

41. S. Koren et al., Canu: scalable and accurate long-read assembly via adaptive k-mer weighting and repeat separation. Genome Res 27, 722–736 (2017).

42. M. Alonge et al., Automated assembly scaffolding using RagTag elevates a new tomato system for high-throughput genome editing. Genome Biol 23, 258 (2022).

43. M. Nattestad, M. C. Schatz, Assemblytics: a web analytics tool for the detection of variants from an assembly. Bioinformatics 32, 3021–3023 (2016).

44. H. Li, Minimap2: pairwise alignment for nucleotide sequences. Bioinformatics 34, 3094–3100 (2018).

45. G. Mohannath, C. S. Pikaard, Analysis of rRNA Gene Methylation in Arabidopsis thaliana by CHEF-Conventional 2D Gel Electrophoresis. Methods Mol Biol 1455, 183–202 (2016).

46. A. Wilm et al., LoFreq: a sequence-quality aware, ultra-sensitive variant caller for uncovering cell-population heterogeneity from high-throughput sequencing datasets. Nucleic Acids Res 40, 11189–11201 (2012).

