## Supplemental information-text and figs for "Sequence and epigenetic landscapes of active and silenced nucleolus organizers in Arabidopsis"

**The PDF file includes:**

Materials and Methods
Figs. S1 to S8
Tables S1 to S2

**Other Supplementary Materials for this manuscript include the following:**

Data S1 to S10

### Materials and Methods

Plant material

*Arabidopsis thaliana* Col-0 (Arabidopsis Biological Resource Center stock CS1092), Col-0 bearing the Fib2:YFP transgene (*40*), and Col-0 Fib2:YFP, *hda6-6* mutants plants (*15*) were the sources of DNA or RNA used in this study. For three week-old seedling tissue, plants were grown on 0.5x Murashige and Skoog medium (MS) plates. For inflorescence apex and leaf tissue, plants were grown in soil.

PacBio SMRT Sequencing

6 g of frozen inflorescence apex tissue was used for CTAB / phenol / chloroform DNA extraction (for details, see DNA extraction section of https://benchling.com/s/prt-lQHskqJ9QFYOJ1MWhMs4?m=slm-3raRnwghjJbspfwq3k9Z). The resulting genomic DNA was digested with *I-Ppo* I to cut 45S rRNA gene arrays into ~10 kbp units. The digested DNA was subjected to electrophoresis using a 0.7% agarose gel and the ~10 kbp band was purified using Qiagen's Qiaex II Gel Extraction Kit. Libraries were prepared from this DNA using Pacific Bioscience's DNA Sequencing Kit 4.0 and sequencing was conducted using a PacBio RS II system.

Analysis of SMRT Sequencing Data to determine IGS VLE types

Raw PacBio subreads were placed into the ccs command of the SMRT tools 9 package <ccs --minReadScore=0.6 --minLength=5000 --maxLength=13000 --minPasses=1 --minPredictedAccuracy=0.85 $input $output> to generate consensus read sequences. These reads were aligned to the 45S IGS reference sequence used in a prior study (*25*) Aligned sequences that showed significant structural differences to the reference were manually annotated to generate new reference sequences representing a broad set of 45S IGS sequence types. Additional IGS sequences were discovered during subsequent ONT sequencing data assembly and added to the set of 45S IGS sequence types.

Ultra-long ONT sequencing

Genomic DNA from Col-0 plants was extracted using Circulomics Nanobind products and protocols (https://www.pacb.com/support/documentation/?fwp_documentation_search=plant&fwp_workflow_step=sample-preparation&fwp_sort=preserve). For each prep, 350-1000 mg of frozen inflorescence apex tissue (unopened flower buds) was processed using the Circulomics' "Nuclei Isolation – TissueRuptor Plant Tissue Protocol" (Document ID: NUC-TRP-001). High molecular weight DNA was then extracted from the resulting nuclei preparations using Circulomics' "Nanobind HMW DNA Extraction – Plant Nuclei Protocol" (Document ID: EXT-PLH-002). Libraries were prepared using 40 ug of resulting DNA (measured using a Qubit fluorometer) and an Ultra-Long DNA Sequencing Kit (SQL-ULK001). Sequencing was performed for 72 hours using either a MinION or GridION sequencer with R9.4.1 flowcells.

Whole genome assembly

Base-calling of raw ONT sequencing data was performed using Guppy 6.1.2 using the psac (plant super accurate) model. Only reads below 100 kbp were retained. Filtlong (https://github.com/rrwick/Filtlong ) was then used to sample reads to ~30X coverage (filtlong --min_length 1000 --mean_q_weight 5 --target_bases 4200000000 $LONG_READ > ${LONG_READ%.*}_filtlong_30Xquality.fq). Filtered reads were then assembled by canu 2.1 (*41*) (/canu-2.1/bin/canu -p $NAME -d $outpath genomeSize=140m -nanopore $infile useGrid=false maxThreads=24 maxMemory=210 overlapper=mhap utgReAlign=true). The resulting assembly was compared against the TAIR10 nuclear genome using RagTag (*42*) and visualized using Assemblytics (*43*).

Ribosomal RNA gene assembly using dot plots

For similarity matrix (dot plot)-based assembly, base-calling of raw ONT sequencing data was performed with Guppy 3.6.1. 45S rRNA gene containing reads were selected by mapping fastq reads to the consensus rRNA gene reference using minimap2 (*44*). Dot plots were generated with the dottup tool (word size =11) of Geneious 9.0.5. and joined based on overlapping, visually distinct patterns.

NOR assembly computational pipeline

Base-calling of raw ONT sequencing data was performed using Guppy 6.1.2 using the psac (plant super accurate) model. Filtlong was used to first select reads longer than 35 kbp. Reads that mapped to the Col-0 45S rRNA gene consensus upon minimap2 alignment were then selected. To retain the longest reads, 45S rRNA gene reads over 275 kbp were saved and set aside, while remaining reads were put through additional selection steps, first splitting reads at locations with no NOR kmers for more than 12 kbp (filtlong -a references.fa --split 12000) and then filtering for high quality, long reads of at least 85 kbp (filtlong --min_length 85000 --min_mean_q 90). This selected, quality controlled 45S rRNA gene read set was then concatenated with the unfiltered >275 kbp 45S rRNA gene read set, resulting in the final NOR assembly read set. The computational assembly involved the creation of two Snakemake-based pipelines (https://snakemake.readthedocs.io/en/stable/), perread_var_call and tagged_longread_assembler. First, the NOR assembly read set was converted into 'VLE sequence' using the perread_var_call. Briefly, the pipeline split reads into segments starting at 25S rRNA sequences. These segments were then mapped, via minimap2, to each known variant sequence for the different VLE regions. The element variant with the highest minimap2 alignment score was then chosen as the 'variant call' for that VLE. This was repeated for each VLE. The resulting variant calls throughout a sequencing read were then collected in order, creating a 'VLE sequence' for that read. Second, VLE sequences were used to generate maps of the NORs. This was done using scripts from the tagged_longread_assembler repository. Starting from the NOR telomeric and centromere-proximal ends, VLE strings with unique patterns (VLE landmarks) were gathered and visualized using the "single-pattern_to_SValignment.py" script. Using these visualizations, additional VLE landmarks were discovered, confirmed by analysis of their coverage, and 'maps' of the distance between VLE landmarks were created. These maps were confirmed using visualizations of alignments of the VLE sequences generated by "locus-patterns_to_SValignment.py" and color coded using "SValign_to_colored-xlsx.py". These visualizations are provided in files S3-S7. This process allowed the linking of VLE landmarks into contigs across the entirety of both NORS. Third, the DNA sequence assembly was performed using the main "tagged_longread_assembler" pipeline. The map files for each contig, combined with the NOR assemblies read set and VLE sequence files, were used as input to generate polished contigs. Briefly, the pipeline searched the VLE sequences for the VLE landmarks of the map. It then inserted a unique DNA sequence tag at each 25S sequence in the original fastq reads, defining where that portion of the read occurs in the NOR. This allowed all read segments from a given read to be placed into a multiple sequence alignment with other reads corresponding to the same position within a NOR. The resulting consensus sequences for each position were then joined into a contig. The tagged reads were then used for several rounds of racon polishing, with the unique sequence tags forcing proper alignment. The pipeline then removed the unique DNA tags, resulting in polished contigs (see the "tagged_longread_assembler" github for further details). Finally, the polished contigs were merged by hand. Low coverage regions at ends of contigs were trimmed. At the NOR ends, where the NORs join either the telomeres or centromere-proximal non-rRNA gene sequences, junction sequences were merged manually. Regions where NOR contigs could not be unambiguously joined due to extensive gene subtype homogeneity (e.g sites denoted by # in Figure 2) were marked by 100 tandem insertions of "N" into the NOR sequence. This resulted in final sequences for *NOR2* and *NOR4* that were then annotated using Geneious Prime and submitted to Genbank.

<u>Analysis of assembled NOR sequences</u>

*NOR2* and *NOR4* fasta files were placed into the "perread_var_call" pipeline, yielding VLE variant calls for the assembly. The script "SV-kmer_homogeneity.py" from "perread_var_call" was used to analyze the homogeneity of the NORs within a sliding window spanning 9 rRNA genes. Rstudio was used to generate color-coded tracks displaying the positions of VLE variant calls and homogeneity scores along the NORs. The script "SV-kmer_dot-plot.py" from "perread_var_call" was used to generate a similarity matrix using VLE sequence rather than DNA sequence. Rstudio was used to display the output. Similarity matrix analyses comparing NOR VLE patterns to a 45S rRNA gene reference VLE pattern was used to visualize the direction and organization of the repeats.

<u>sgRNAs for targeted Cas9 cutting in the 3' ETS region of different rRNA gene subtypes</u>

We used a custom script to identify potential target sequences present in the 3'ETS variable region, following guidelines for sgRNA design (thttp://www.clontech.com/sgRNA-design-tools). A set of forward PCR primers were then made that included T7 promoter sequences adjacent to the sgRNA target sequences and the Guide-it Scaffold Template-specific sequence for sgRNA synthesis using a Guide-it sgRNA In Vitro Transcription Kit (Cat. No. 632635). RNA products were purified using a Guide-it IVT RNA Clean-Up Kit (Cat. No. 632638). The efficacy of the

different sgRNAs was then tested using a Guide-it sgRNA Screening Kit (Cat. No. 632639). Briefly, genomic DNA (50 ng) purified from *Arabidopsis thaliana* Col-0 plants was incubated with sgRNA (25 ng) in the presence of 250 ng of Cas9 nuclease. The Cas9 cleavage efficiency within the 3'ETS region was then assessed by PCR amplification using a pair of primers flanking the 3'ETS region (F: GACAGACTTGTCCAAAACGCCCACC; R: CTGGTCGAGCTAATCCTGGACGATT). Reactions were then subjected to agarose gel electrophoresis and SYBR Safe staining. The sgRNAs ultimately used in the study (see fig. S4), and their 3' ETS VLE specificities, were
VAR 1+2-targeting sgRNA, ATGAAAACTGGTGATTGTTGCGG;
VAR 1+3-targeting sgRNA, AGAAACGGAAGAGAAAGCGTGGG; and
VAR 1, 3, 4-targeting sgRNA, GAACTAGCAAGTAATCGTCCAGG.

Analysis of genomic sgRNA-CAS9 digestion by CHEF gel electrophoresis and Southern blotting

Ultra-high molecular weight DNA was prepared as described for ONT sequencing and embedded in agarose plugs. Agarose plugs were then placed in 50 mL conical tubes and incubated in 10 mL of T10E10 buffer (10 mM Tris-HCl, 10 mM EDTA, pH 8.0) supplemented with 2 mM PMSF for 1 hr at 4°C. Plugs were then washed four times, 30 min each, at room temperature in 10 mL of T10E10 buffer without PMSF. Next, individual agarose plugs were washed twice, 1 hr, at room temperature, with 1 ml of 1× Cas9 reaction buffer (NEB, cat. #. B0386). After a second wash, the plug was incubated for 1 hr, at 37°C, with 100 µL of 1× Cas9 reaction buffer (NEB) containing 200 µM of sgRNAs and 1 µM Cas9 enzyme. The buffer was then removed and replaced with 500 µL of 20 mM Tris-HCl, 50 mM EDTA, pH 8.0 and incubated at 10 min at room temperature. The agarose plug was then subjected to 5 washes, each 15 min at room temperature, with 10 mL of TE buffer. Agarose plugs were inserted into the wells of a precast 1% Certified Megabase Agarose (Bio-Rad) gel and subjected to CHEF electrophoresis using a Bio-Rad sytem. Running parameters for stage I were: initial and final switch time of 60 sec, 200V, run time 13 hr and for stage II were: initial and final switch time of 90 sec, 200V, Run time 7 hr. The gel was then subjected to Southern blotting as described by Mohannath and Pikaard (*45*). The digoxigenin (Dig)-labeled 25S rDNA probe was synthesized using a PCR Dig Probe Synthesis kit (Roche, 11636090910) and primers CCGGAGGTAGGGTCCAGCGG (forward) and CCGCCGTTTACCCGCGCTTG (reverse). Hybridization was performed at 42°C, for 72 hr, followed by membrane washing and blocking using DIG Wash and Block Buffer Set (Roche, 11585762001). rRNA gene fragments were detected using anti-Dig AP antibodies (1:15,000 dilution) (Roche, 11093274910) and CDP-Star Detection (Roche, 11759051001).

Whole genome Illumina sequencing

Genomic DNA (gDNA) was isolated from Arabidopsis seedlings using the Qiagen DNeasy Plant Mini Kit (Cat No: 69106). The gDNA library was prepared using the Illumina Nextera Flex kit (Cat No: 20025523) and then sequenced on an Illumina NextSeq instrument to obtain 75 bp paired-end reads. Resulting sequence data was subjected to quality control and trimming using

Trimmomatic (https://github.com/usadellab/Trimmomatic). The data was then mapped to a 45S rDNA consensus sequence using Bowtie2, with allowance for a single mismatch in the seed region. The mapped reads were variant-called using LoFreq (*46*)(https://csb5.github.io/lofreq/) which generated a VCF with all the variant calls across a single rDNA unit. The --no-default-filter flag was used to run LoFreq with high sensitivity and then filtered for sites predicted to occur with a frequency higher than .001. The resulting VCF file was input into the custom snakemake pipeline "perread_SNP_call" along with the NOR assembly sequences. Briefly, the first portion of this pipeline maps each unit of the input sequences (NOR assembly) to the reference (45S reference), then looks at the mapped bases at each position of interest (as defined by the LoFreq vcf), outputting a line of variant calls for each unit. Rstudio was used to generate a heatmap display of reference versus variant nucleotides positioned along the NORs.

<u>RNA sequencing</u>

Mature leaf of three-week-old plants or inflorescence tissue was collected, and total RNA was extracted using a Zymo Research Quick-RNA Plant Miniprep Kit (Cat No: 11-362). The resulting RNA was treated with Turbo DNase (Cat No: AM2238). The library was prepared using an Illumina TruSeq Stranded total RNA HT kit with no enrichment or depletion steps. The library was sequenced on an Illumina NextSeq instrument to obtain 75 bp paired-end reads. Resulting sequence data were subjected to quality control and trimmed using Trimmomatic. The data were mapped to 45S rDNA consensus sequence using Bowtie2, allowing a single mismatch in the seed region. The mapped reads were then filtered for those that mapped in the forward direction and these were then variant-called using LoFreq with the --no-default-filter flag to maximize sensitivity. Resulting VCF files were input into the second portion of the "perread_SNP_call" pipeline. This takes the alternate allele frequency from the RNA-seq vcf, normalized the values based on the number of occurrences of that allele in the assembly, and then output the normalized values at the location of the variant allele (polymorphism relative to the reference sequence) on the NORs. The resulting normalized expression values were displayed in a heatmap using Rstudio.

<u>Flow-sorting of nuclei and nucleoli, DNA isolation and sequencing</u>

To purify nuclei, ~ 2 g of *Arabidopsis thaliana* leaf tissue from ~ 3 week old seedlings (wild-type or *hda6* mutant Col-0 expressing Fib2:YFP) were fixed for 20 min in 4% formaldehyde in Tris buffer (10 mM Tris-HCl [pH 7.5], 10 mM EDTA and 100 mM NaCl), washed twice for 10 min with ice-cold Tris buffer, and minced with a razor blade in 1 mL of 45 mM MgCl2, 20 mM MOPS [pH 7.0], 30 mM sodium citrate, and 0.1% Triton X-100. The homogenate was filtered through a 30 µm mesh (Sysmex CellTrics) and subjected to fluorescence activated nuclear sorting (FANS), triggered by the Fib2:YFP signal, using a BD FACS Aria II instrument as described previously (*15*). To purify nucleoli, the homogenate was subjected to sonication with a Diagenode Bioruptor (three 5-min pulses; medium power), which disrupts nuclei. The sonicated material was then subjected to fluorescence-activated nucleolar sorting (FANoS) (*15*). Approximately 1 million sorted nuclei and ~ 1.5 million sorted nucleoli were collected. The nuclei and nucleoli samples were then treated with Proteinase K (100 µL of 20 mg/mL stock solution/1 mL of sample) and RNase A (20 µL of 10 mg/mL stock solution /1mL of sample). After gentle mixing by tube inversion, the samples were incubated at 50˚C for 30 min, followed by incubation at 95˚C for 20 min. 250 µL of 4x DNA extraction buffer (0.55 M Tris-HCl [pH 8.0], 1 M NaCl, 100 mM EDTA) was then added per 1 mL of final sample volume and mixed gently by tube inversion. DNA was precipitated by the addition of 1 volume of ice-cold isopropanol and incubation on ice for 1 hr in the presence of 1 µL (15 µg/µL) of Glycoblue. DNA was recovered by centrifugation of 16,000g for 15 min at 4C. The DNA pellet was washed twice with 1 mL of ice-cold 70% ethanol and resuspended in 1x TE buffer. DNA libraries were prepared using an ONT Rapid Sequencing Kit (SQK-RAD004) and sequenced using a GridION sequencer with R9.4.1 flowcells.

Analysis of 5-methylcytosine (5mC) positions in ONT sequencing data

Detection of 5mC modification was performed using deepsignal-plant (*30*), as described in the 'Usage' section of the associated Github (https://github.com/PengNi/deepsignal-plant), using raw sequencing data (fast5s) corresponding to the fastq reads used for the NOR assemblies. This was followed by quantification of methylcytosine frequency according to sequence context (CG, CHG or CHH) using the included python script. The resulting 5mC frequencies for each cytosine in the NOR assemblies were binned using sliding windows of 500bp, with a 250bp step. Resulting values were then imaged in xy plots using Rstudio.

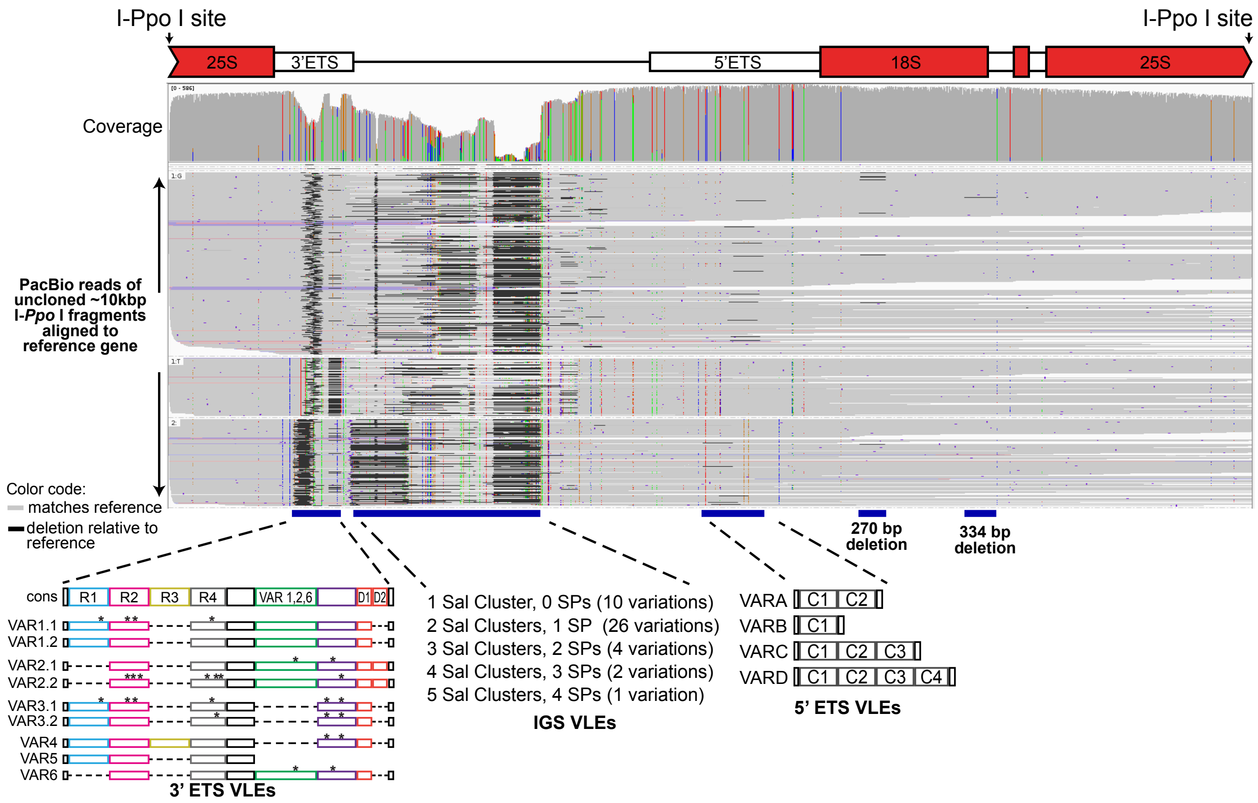

**Fig. S1. Variation identified by PacBio sequencing of rRNA gene repeat units.**
Genomic DNA was digested with the rDNA-specific endonuclease *I-Ppo* and resulting ~10 kb fragments were gel-purified and deep-sequenced. Individual reads were then aligned to an rRNA gene reference sequence (see file S1). Sequences identical to the reference are shown in gray. Differences relative to the reference are shown in black or other colors. Intervals corresponding to the VLEs used to define rRNA gene subtypes are shown at the bottom of the figure, as in Figure 1B.

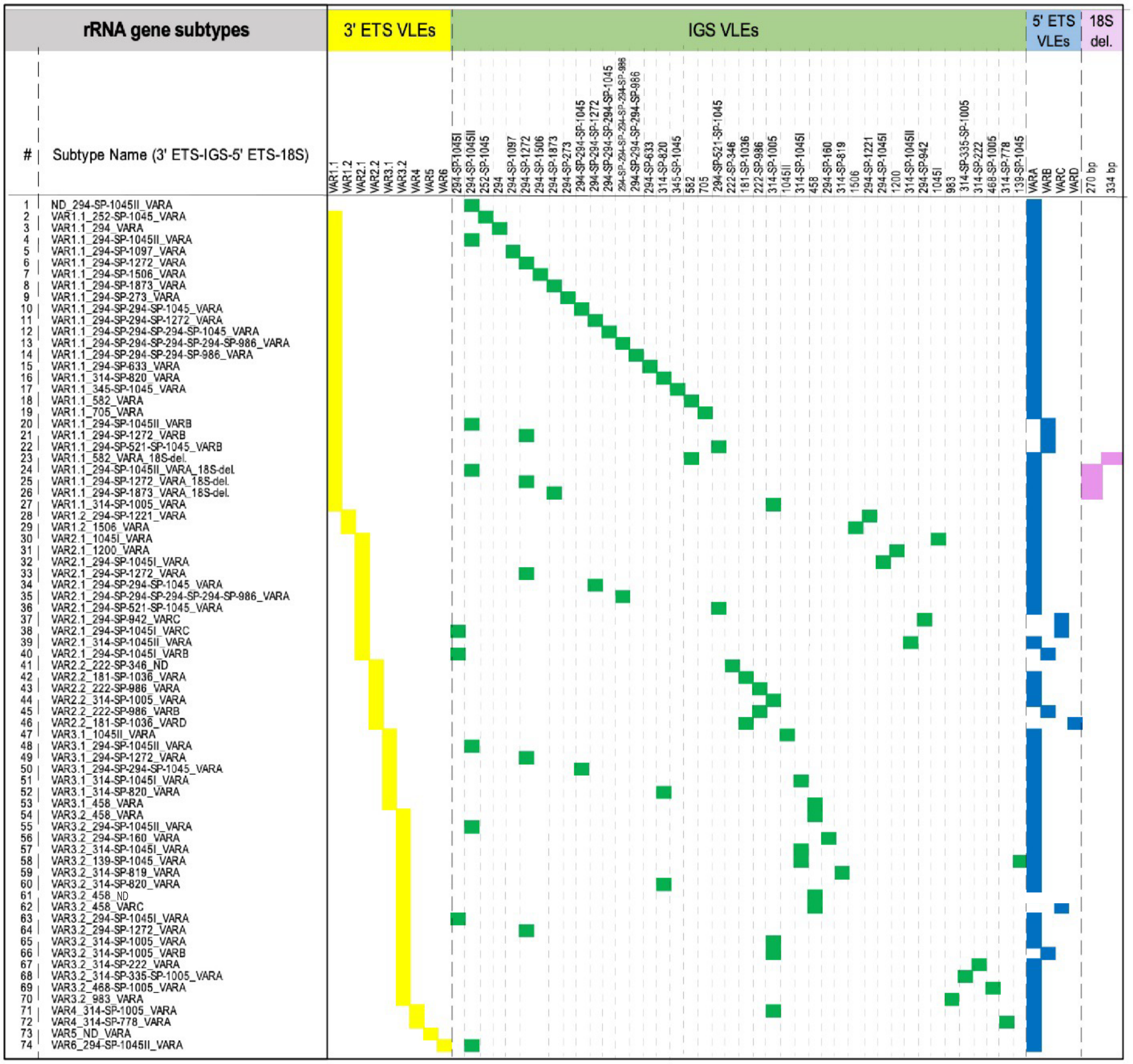

**Fig. S2A. Gene subtypes defined by their VLE compositions.**

The 59 VLEs summarized in Figure 1B occur in 74 different combinations among rRNA genes, thus defining 74 distinct rRNA gene subtypes. The most prevalent VLE is VARA, occurring in 64 gene subtypes and corresponding to the presence of two tandem C repeats within the 5' ETS. VLEs corresponding to different 3' ETS sequences provide the next largest groupings, with 27 gene subtypes having VAR1-class VLEs (with point mutations allowing 1.1 or 1.2 sub-classifications), twenty-four having VAR3.1 or 3.2 VLEs and nineteen having VAR2.1 or 2.2 VLEs. The least frequent subtypes carry VAR4, VAR5, and VAR6 VLEs. Subtype-specific VLEs occur primarily in the IGS, where there can be as few as one, or as many as five, clusters of Sal repeats (each repeat being 20-21 nt long) that vary in number. Sal repeat cluster lengths, in bp, are given in the gene subtype names. Likewise, variable numbers of spacer promoters (SPs) occur when there are two or more Sal repeat clusters. In total, there are 42 different Sal cluster-SP arrangements, 27 of which are unique to single rRNA gene subtypes. The remaining 15 IGS arrangements occur in two or more subtypes, an example being 294-SP-1272, having Sal repeat clusters of 294 bp and 1272 bp separated by a single spacer promoter. This IGS arrangement is present in six subtypes that collectively represent the 3 major 3'ETS classes, VAR1, VAR2 and VAR3.

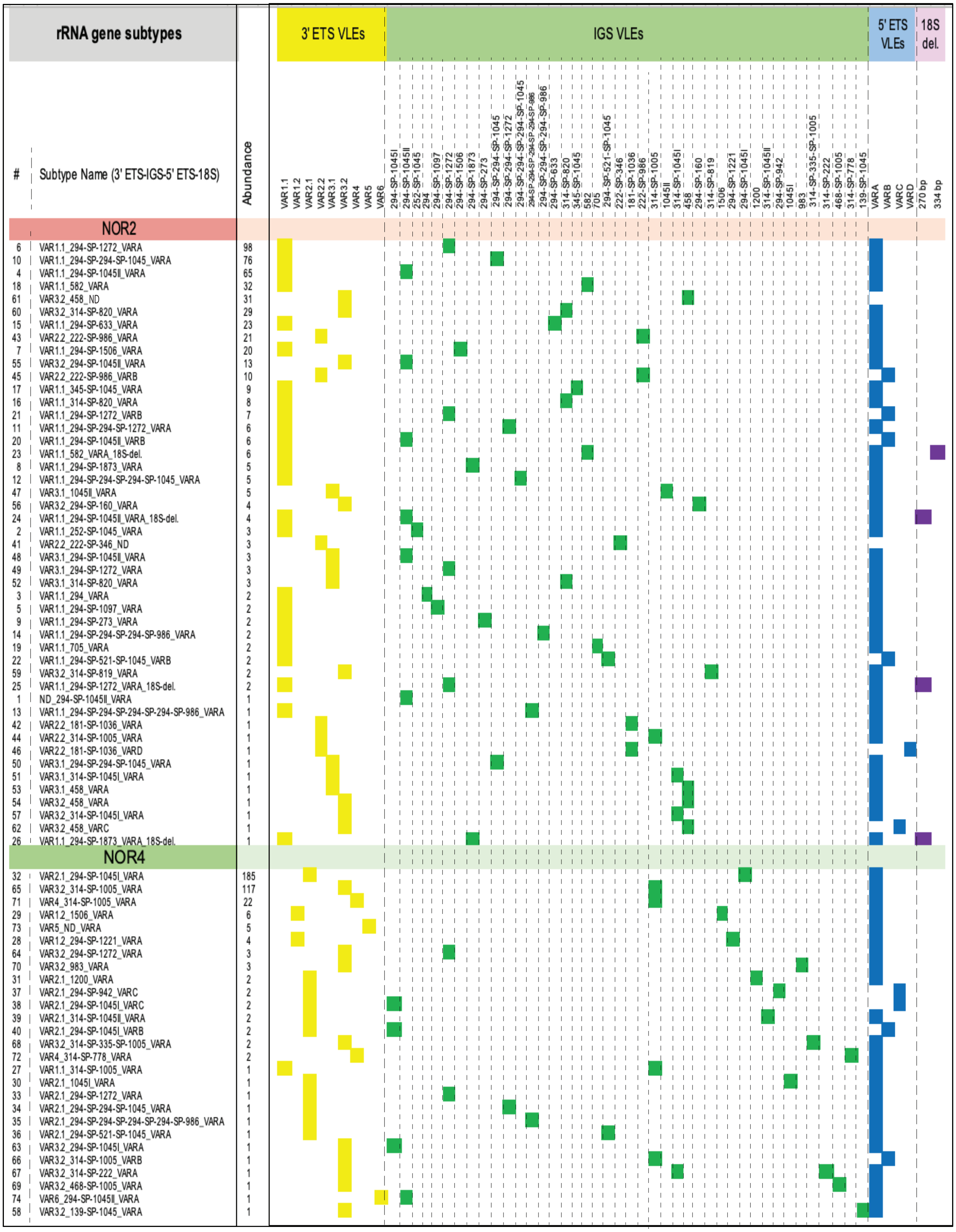

**Fig. S2B. rRNA gene subtypes sorted according to *NOR2* or *NOR4* affiliation.**

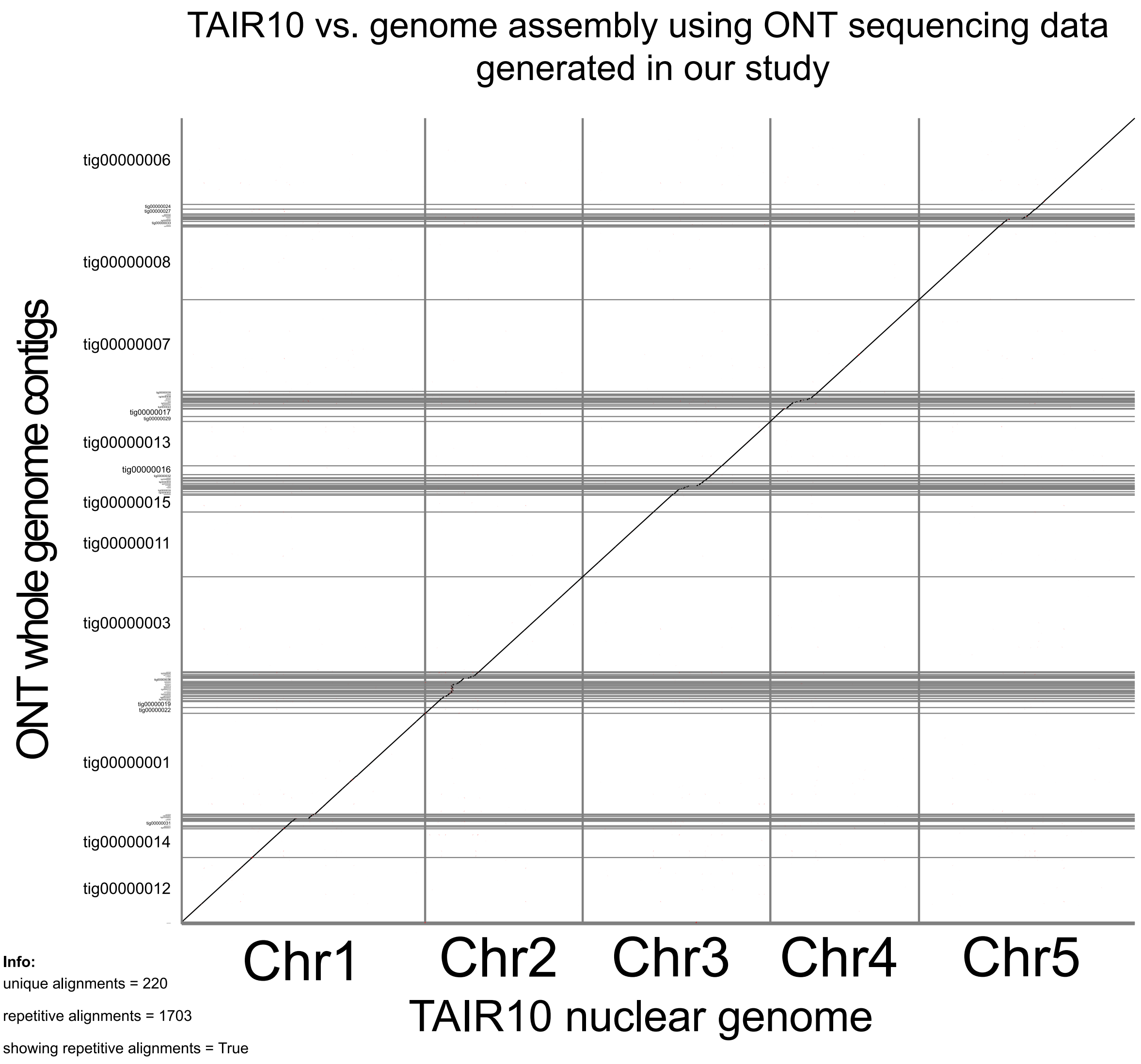

**Fig. S3. Chromosome sequences assembled using ONT reads of the current study are contiguous with those of the TAIR10 assembly.**

Contigs assembled using ultra-long ONT reads and Canu assembly tools (see methods) were scaffolded to the Arabidopsis nuclear chromosome assemblies from TAIR10 using RagTag. The similarity matrix was generated using the Assemblytics tool. The ONT sequence assembly aligns with the majority of the TAIR10 reference with no indicators of large rearrangements. Observed discontinuities occur in centromeric regions.

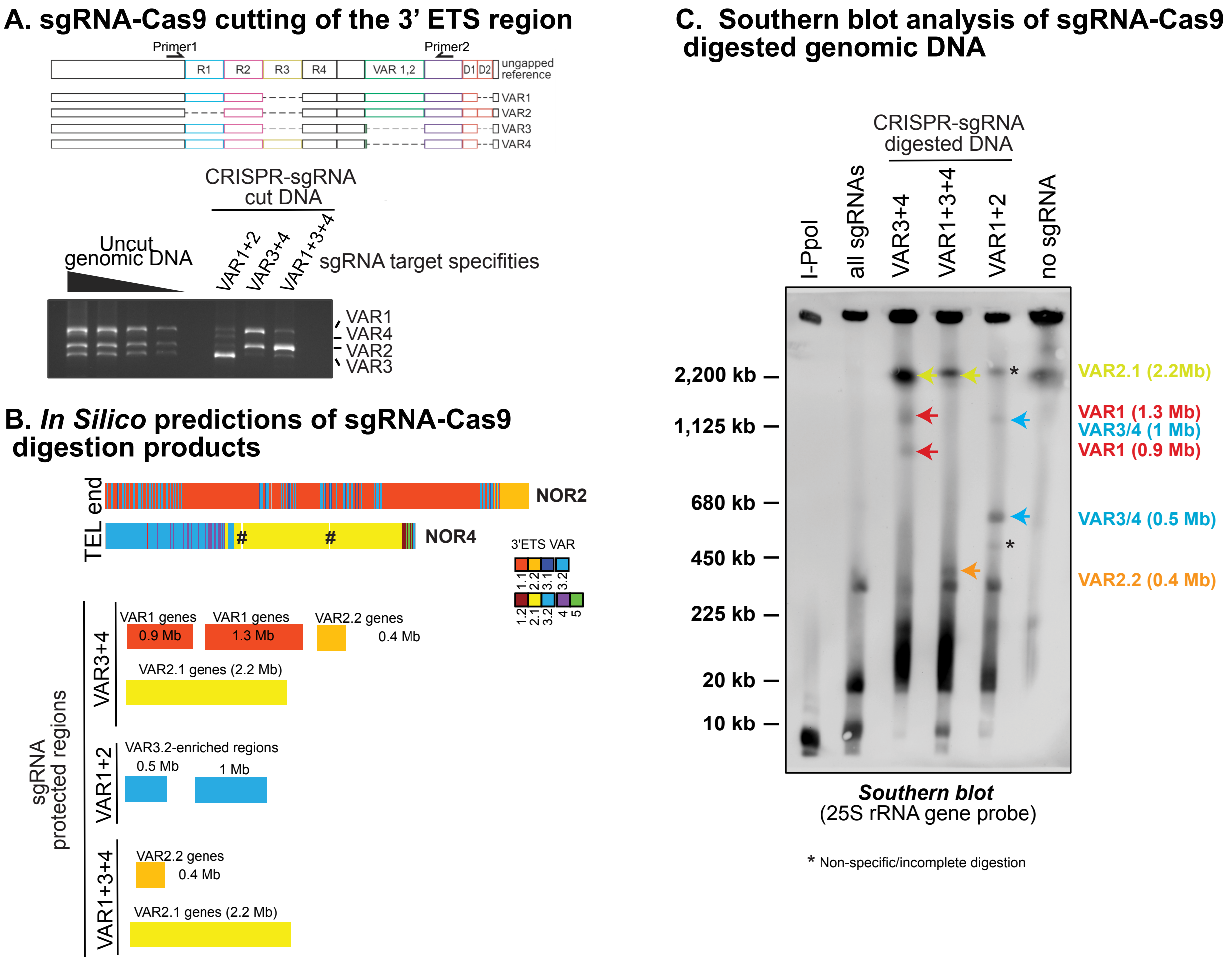

**Fig. S4. Physical mapping test of the NOR assemblies using custom guide RNAs directing Cas9 cleavage of rRNA gene 3’ ETS regions**

(**A**) The diagram shows the 3’ETS variable region and the positions of PCR primers that flank the region. PCR amplification of genomic DNA using these primers yields products of different lengths, corresponding to the abundant genes bearing the VAR1, VAR2 and VAR3 VLEs and the less abundant genes bearing the VAR4 VLE. The stained agarose gel shows the amplification products obtained using either uncut genomic DNA (left lanes) or DNA that had been incubated with Cas9 and three different sgRNAs prior to PCR. Two of the sgRNAs guide the digestion of genes of two different VLE classes that share the same sgRNA target sequence (either VAR1 + VAR2 or VAR3 + VAR4), and the third sgRNA targets three VLE classes (VAR1 + VAR3 + VAR4). Note that the targeted VLE classes are depleted among the PCR amplification products, demonstrating the specificity and efficacy of the sgRNA-Cas9 complexes. (**B**) *In silico* prediction of large sgRNA-Cas9 digestion fragments of *NOR2* and *NOR4* based on the sgRNA specificities demonstrated in panel A. The sizes of the expected fragments are shown, with color-coding showing the regions of the NORs giving rise to the fragments. (**C**) sgRNA-Cas9 digestion products visualized by CHEF gel electrophoresis and Southern blotting with a 25S rRNA probe. For this experiment, ultra-high molecular weight gDNA was embedded in agarose plugs and subjected to Cas9 digestion programmed by individual sgRNAs, as in panel A, or a mix of all three sgRNAs. I-*Ppo*I and no-digestion controls are included in the first and last lanes. The DNA fragments were resolved by CHEF electrophoresis and visualized by Southern blotting and hybridization to the 25S rDNA probe. Predicted large fragments (see panel B) were observed.

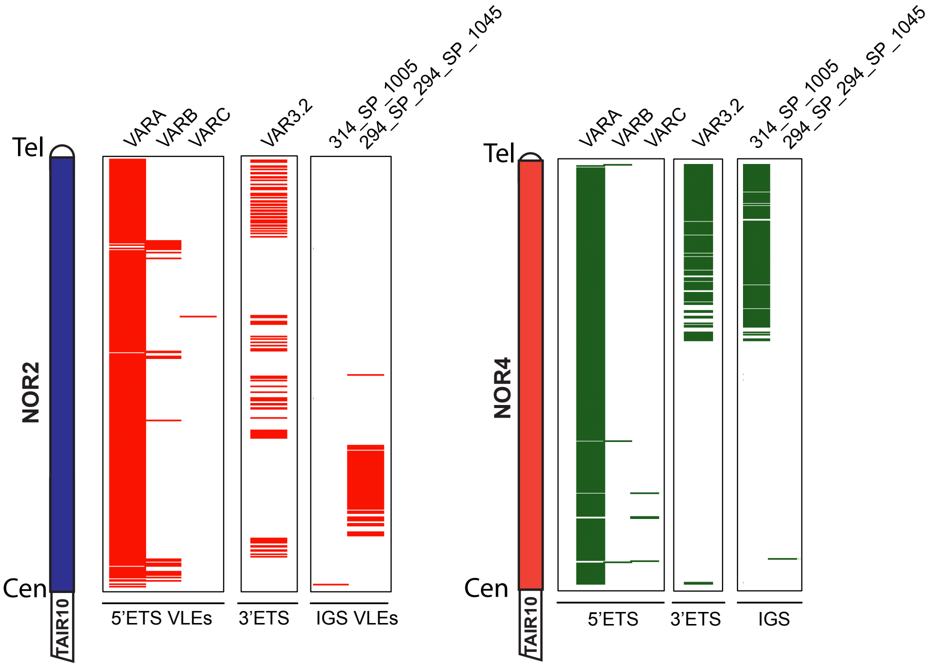

**Fig. S5. VLEs that are common to both NORs.**

The positions of 6 VLEs present within genes of both *NOR2* and *NOR4* are indicated by colored horizontal lines. Tel and Cen indicate the telomere and centromere-proximal ends of the NORs. TAIR10 indicates where the current sequences of chromosomes 2 and 4 begin in the TAIR10 genome assembly.

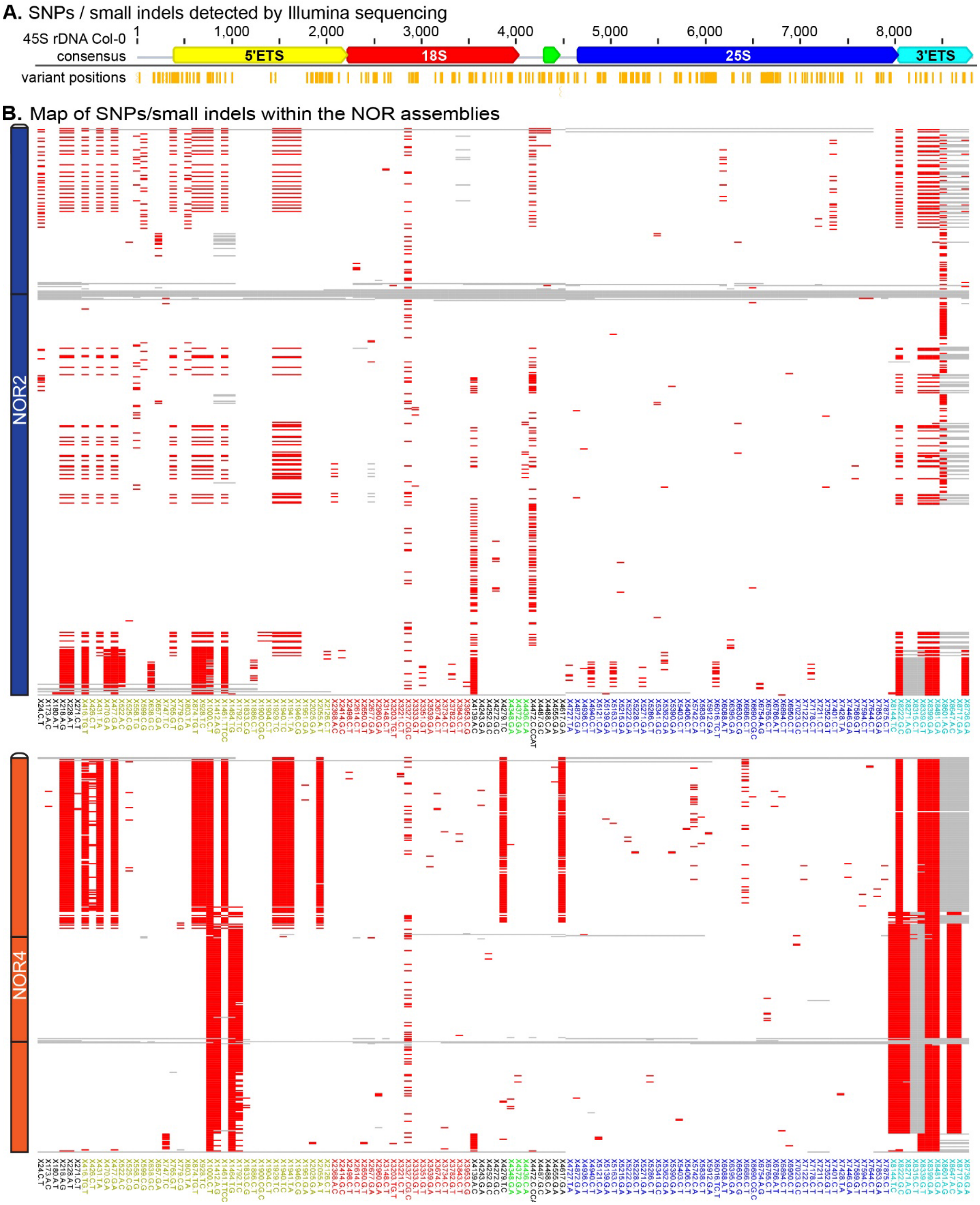

**Fig. S6.** Illumina sequencing-detected SNPs/small indel positions within individual genes of *NOR2* and *NOR4*. (**A**) Vertical orange bars below the rRNA gene diagram represent positions of SNPs / small indels (relative to the reference gene) detected in Illumina sequencing reads using LoFreq. (**B**) Map of SNPs/small indels on the NOR assemblies. Each line on the Y axis represents an individual 45S gene, from positions near the telomere (top) to centromere (bottom). Each column marks the presence (red) or absence (white) of SNPS/small indels. Gray represents no alignment to the consensus at that region. SNP labels on the x-axis are color-coded to match the regions in (A).

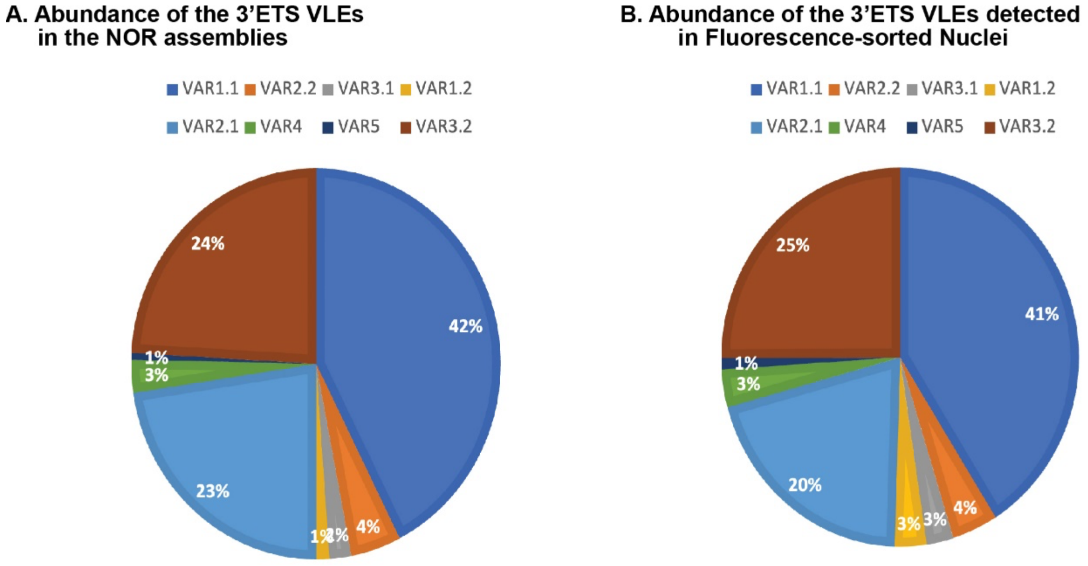

**Fig. S7. rRNA gene subtype abundance in flow-sorted whole nuclei closely matches their abundance in the NOR sequence assemblies.**

(**A**) Abundance of genes bearing the different 3’ETS VLEs in the NOR assemblies versus (**B**) the abundance of the genes bearing the different 3’ETS VLEs following their detection by ONT sequencing of DNA purified from flow-sorted nuclei. This experiment indicates that flow-sorting allows all subtypes to be detected without apparent bias.

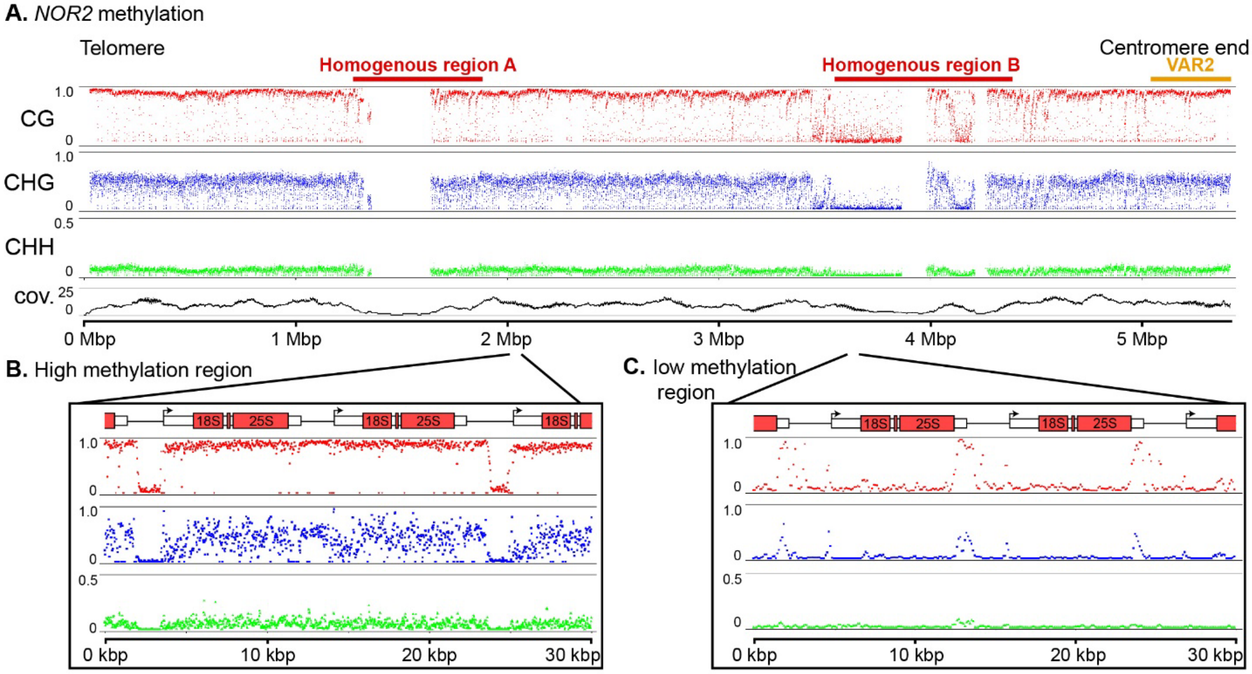

### Fig. S8. Details of rRNA gene methylation patterns.

(**A**) 5mC frequencies in the CG, CHG, and CHH contexts at *NOR2* are shown, as in Figure 5A. (**B**) Zoomed-in view of a region characterized by high 5mC levels showing the dips in methylation that occur upstream of many, but not all, gene promoter regions. (**C**) Zoomed-in view of a representative region characterized by low 5mC levels, showing the characteristic spike in methylation observed at the 3'ETS region.

### Table S1. ONT sequencing run statistics

#### Total output

| ONT Sequencing Run | Run Name | # cells | total bp: | | total bp per flowcell: | |
|---|---|---|---|---|---|---|
| | | | 100kb+ | 200kb+ | 100kb+ | 200kb+ |
| 2022-05-12 Col-0 | 2022_5_12_CS1092_floapex_nuclei_ULK | 2 | 6,020,711,635 | 1,561,915,198 | 3,010,355,818 | 780,957,599 |
| 2022-07-28 Col-0 | 2022_07_28_CS1092_infloapex_ULK | 5 | 6,831,857,395 | 1,616,646,803 | 1,366,371,479 | 323,329,361 |
| Combined runs | | 7 | 12,852,569,030 | 3,178,562,001 | | |

#### 45S gene output

| ONT Sequencing Run | Run Name | # cells | 45S rRNA gene total bp: | | 45S rRNA gene bp per flowcell: | |
|---|---|---|---|---|---|---|
| | | | 100kb+ | 200kb+ | 100kb+ | 200kb+ |
| 2022-05-12 Col-0 | 2022_5_12_CS1092_floapex_nuclei_ULK | 2 | 456,785,638 | 128,525,522 | 228,392,819 | 64,262,761 |
| 2022-07-28 Col-0 | 2022_07_28_CS1092_infloapex_ULK | 5 | 581,480,077 | 157,498,739 | 116,296,015 | 31,499,748 |
| Combined runs | | 7 | 1,038,265,715 | 286,024,261 | | |

**bp in quality-controlled assembly read set:**

931,988,411

### Table S2.

**Sequencing data statistics for fluorescense-sorted nuclei and nucleoli**

| | Total reads | Total DNA bases | Read Length (N50) | 45S ribosomal DNA reads | 45S ribosomal DNA bases | 45S ribosomal DNA Read Length (N50) |
|---|---|---|---|---|---|---|
| WT Nuclei | 869,619 | 1,428,395,669 | 3,180 | 84,472 | 185,912,945 (13% of Total) | 4,285 |
| WT Nucleoli | 77,876 | 113,550,068 | 2,615 | 33,713 | 57,184,319 (50% of Total) | 2,864 |
| *hda6* Nuclei | 527,843 | 874,477,467 | 3,569 | 23,204 | 73,745,623 (8.4% of Total) | 6,797 |
| *hda6* Nucleoli | 48,148 | 91,843,471 | 4,627 | 12,357 | 35,787,141 (39% of Total) | 5,619 |

**Data S1. (separate file)**

45S rRNA gene consensus reference sequence (fasta file) used for VLE analyses and sequence for subtype #10, used for dot-plot analyses of ONT reads

**Data S2. (separate file)**

NOR assembly landmarks, observed vs. predicted coverage based on sequencing depth.

**Data S3. (separate file)**

ONT read alignments based on VLEs for *NOR2* telomere end.

**Data S4. (separate file)**

ONT read alignments based on VLEs for *NOR2* centromere end.

**Data S5. (separate file)**

ONT read alignments based on VLEs for *NOR4* telomere end.

**Data S6. (separate file)**

ONT read alignments based on VLEs for *NOR4* central region.

**Data S7. (separate file)**

ONT read alignments based on VLEs for *NOR4* centromere end.

**Data S8. (separate file)**

Nucleotide accuracy for NOR assemblies.

**Data S9. (separate file)**

NOR assemblies capture the VLE content of the sequencing reads.

**Data S10. (separate file)**

The NOR assemblies represent the variation detected in Illumina sequencing.
